# Neuronal avalanches in input and associative layers of auditory cortex

**DOI:** 10.1101/620781

**Authors:** Zac Bowen, Daniel E. Winkowski, Saurav Seshadri, Dietmar Plenz, Patrick O. Kanold

## Abstract

The primary auditory cortex processes acoustic sequences for the perception of behaviorally meaningful sounds such as speech. Sound information arrives at its input layer 4 from where activity propagates to associative layer 2/3. It is currently not known whether there is a particular organization of neuronal population activity that is stable across layers and sound levels during sound processing. We used *in vivo* 2-photon imaging of pyramidal neurons in cortical layers L4 and L2/3 of mouse A1 to characterize the populations of neurons that were active spontaneously, i.e. in the absence of a sound stimulus, and those recruited by single-frequency tonal stimuli at different sound levels. Single-frequency sounds recruited neurons of widely ranging frequency selectivity in both layers. We defined neural ensembles as neurons being active within or during successive temporal windows at the temporal resolution of our imaging. For both layers, neuronal ensembles were highly variable in size during spontaneous activity as well as during sound presentation. Ensemble sizes distributed according to power laws, the hallmark of neuronal avalanches, and were similar across sound levels. Avalanches activated by sound were composed of neurons with diverse tuning preference, yet with selectivity independent of avalanche size. Thus, spontaneous and evoked activity in both L4 and L2/3 of A1 are composed of neuronal avalanches with similar power law relationships. Our results demonstrate network principles linked to maximal dynamic range, optimal information transfer and matching complexity between L4 and L2/3 to shape population activity in auditory cortex.

## Introduction

The primary auditory cortex (A1) is central to encoding sound sequences (Nelken et al., 2003; Bartho et al., 2009; Francis et al., 2018), yet several major aspects of sound processing in A1 are currently not well understood. First, the selectivity of single A1 neurons to stimulus features such as sound frequency changes with sound level, i.e. across the dynamic range, while perceptual discrimination performance is largely level-invariant (Jesteadt et al., 1977; Wier et al., 1977; Bernstein and Oxenham, 2006). Second, responses of A1 neurons can be unreliable over repeated trials despite robust behavioral outcome (Deweese and Zador, 2004). It was recently shown that auditory stimuli are encoded by populations of A1 neurons (Bathellier et al., 2012; Francis et al., 2018). Here we sought to identify characteristics of active populations in A1.

Identifying population dynamics central to sound processing needs to take into account that A1 is composed of several cortical layers each transforming incoming information (Atencio and Schreiner, 2009; Atencio et al., 2009). Specifically, sensory inputs arriving in layer 4 (L4) are relayed to layers 2/3 (L2/3) via divergent and heterogeneous feed-forward projections (Meng et al., 2017). In this hierarchy, it is not well understood how population dynamics in L4 contributes to neuronal responses in L2/3 raising the general question of how neural representations of sensory stimuli are reliably transmitted through multilayer networks. At least two lines of evidence suggest that ensemble dynamics could support level-independent reliable sound encoding across networks of neurons. First, network dynamics that support a large dynamic range, i.e. that differentiate sounds at both high and low levels, should be of particular interest to auditory processing where sound amplitude can vary over many orders of magnitude. Simulations (Kinouchi and Copelli, 2006; Shriki and Yellin, 2016) and experiments *in vivo* (Gautam et al., 2015) and *in vitro* (Shew et al., 2009; Shew et al., 2010; Shew et al., 2011; Shew and Plenz, 2013) have shown that cortical networks that display neuronal avalanches (Beggs and Plenz, 2003; Petermann et al., 2009; Shriki et al., 2013; Bellay et al., 2015) maximize their dynamic range for input processing as well as their information capacity (for review see (Chialvo, 2010; Shew and Plenz, 2013)). Neuronal avalanches are identified by power law relationships for the size and duration of activity in neuronal groups which naturally links to a second line of evidence identifying conditions for which information transmission between complex networks is maximized. Theory suggests that complex networks exchange maximal information if their complexity, quantified by their respective power law relationships, is matched (West et al., 2008; Aquino et al., 2010; Aquino et al., 2011). Accordingly, we hypothesize that if A1 population dynamics are governed by avalanche statistics, these statistics would be similar for L4 and L2/3.

Here, using *in vivo* 2-photon imaging of pyramidal neurons in mouse A1, we identify common dynamical principles in sound encoding for both layers that allow for the encoding of sounds in neuronal populations in line with predictions for neuronal avalanches.

## Methods

All procedures were approved by the University of Maryland Institutional Animal Care and Use Committee. Mice were housed under a reversed 12 h-light/12 h-dark light cycle with ad libitum access to food and water. Imaging experiments were generally performed near the end of the light and beginning of the dark cycle. Age range at the time of experiments extended from P46-P100 (75 ± 21 days, mean ± std). Mice of both sexes were used (n=12, 8 Male, 4 Female) based on their availability not by any biased selection.

### Animals

Mice used for this study expressed the genetically encoded calcium indicator (GCaMP6s) (Chen et al., 2013). GCaMP6s was introduced in one of the following ways. In a first line of experiments, we injected an adeno-associated virus (AAV1) into primary auditory cortex (A1) of C57/BL6 mice (n=5) for delivery of a calcium indicator with co-expression of a structural marker mRuby2 (Rose et al., 2016). Both proteins were under control of the synapsin promoter. Virus (AAV1.hSyn1.mRuby2.GSG.P2A.GCaMP6s.WPRE.SV40; titer: 3×10^13^) was obtained from the University of Pennsylvania Vector Core. Observed expression in these mice was primarily limited to supragranular and infragranular layers and was mostly absent in L4. Only data acquired from L2/3 in these animals was included. In a second line of experiments, we used transgenic mice that expressed GCaMP6s either under the Thy1 promoter (Dana et al., 2014, Jax: 024274, GP4.3) or a transgenic mouse line that conditionally expresses GCaMP6s when crossed to a mouse driver line expressing Cre recombinase under the control of Emx1 (Gorski et al., 2002; Madisen et al., 2015, GCaMP6s mouse: Jax: 024115; Emx1-Cre mouse: Jax: 005628). Both L2/3 and L4 datasets were acquired from the transgenic mice. The transgenic L2/3 and L4 data sets were always obtained sequentially in the same mouse, meaning the experiment (sound presentations) was conducted in one layer and then sequentially conducted in the other layer. The two transgenic lines exhibited robust indicator expression in both L4 and L2/3. We found no significant differences of indicator expression within layers when we compared across transgenic mouse lines and animals with viral expression and thus combined the data from different sources according to laminar position.

### Surgery and animal preparation

Mice were given a subcutaneous injection of dexamethasone (5mg/kg) at least 2 hours prior to surgery to reduce potential inflammation and edema from surgery. Mice were deeply anesthetized using isoflurane (5% induction, 2% for maintenance) and given subcutaneous injections of atropine (0.2 mg/kg) and cefazolin (500 mg/kg). Internal body temperature was maintained at 37.5 °C using a feedback-controlled heating blanket. The scalp fur was trimmed using scissors and any remaining fur was removed using Nair. The scalp was disinfected with alternating swabs of 70% ethanol and betadine. A patch of skin was removed, the underlying bone was cleared of connective tissue using a bone curette, the temporal muscle was detached from the skull and pushed aside, and the skull was thoroughly cleaned and dried. A thin layer of cyanoacrylate glue (VetBond) adhesive was applied to the exposed skull surface and a custom machined titanium head plate (based on the design described in Guo et al. (2014) was affixed to the skull overlying the auditory cortex using VetBond followed by dental acrylic (C&B Metabond). A circular craniotomy (~3 mm diameter) was made in the center opening of the head plate and the patch of bone was removed. For wild-type mice, virus (AAV1-syn-mRuby2-GC6s) was loaded into beveled glass pipettes and injected slowly into the areas corresponding to primary auditory cortex in 3-5 sites (~30 nL/site; ~250 – 300 μm from the surface; ~2 – 3 minutes/each injection site). Pipettes were left in place for at least 5 minutes after completion of each injection to prevent backflow. Then, a chronic imaging window was implanted. The window consisted of a stack of 2 – 3 mm diameter coverslips glued with optical adhesive (Norland 71, Edmund Optics) to a 5 mm diameter coverslip. The edges of the window between the glass and the skull were sealed with a silicone elastomer (Kwik-Sil) and then covered with dental acrylic. The entire implant except for the imaging window was then coated with black dental cement created by mixing standard white powder (Dentsply) with iron oxide powder (AlphaChemical, 3:1 ratio) (Goldey et al., 2014). Meloxicam (0.5 mg/kg) and a supplemental dose of dexamethasone were provided subcutaneously as a post-operative analgesic. Animals were allowed to recover for at least 1 week prior to the beginning of experiments.

### Acoustic stimulation

Sound stimuli were synthesized in MATLAB using custom software (courtesy of P. Watkins, UMD), passed through a multifunction processor (RX6, TDT), attenuated (PA5, Programmable Attenuator), and delivered via ES1 speaker placed ~5 cm directly in front of the mouse. The sound system was calibrated between 2.5 and 80 kHz and showed a flat (±3 dB) spectrum over this range. Overall sound pressure level (SPL) at 0 dB attenuation was ~90 dB SPL (for tones). Sounds were played at three sound levels (40, 60, and 80 dB SPL). Auditory stimuli consisted of sinusoidal amplitude-modulated (SAM) tones (20 Hz modulation, cosine phase), ranging from 3 – 48 kHz. For wide-field imaging, the frequency resolution of the stimuli was 1 tone/octave; for 2-photon imaging, the frequency resolution was 2 tones/octave (0.5 octave spacing). Each of these tonal stimuli was repeated 5 times with a 4 – 6 s interstimulus interval for a total of either 75 (wide-field) or 135 (2-photon) iterations.

### Wide-field imaging

In order to construct sound-evoked response maps using wide-field imaging, awake mice were placed into a plastic tube with a head restraint system similar to that described by Guo et al. (2014). Blue excitation light was provided by an LED (470 nm, Thorlabs) or xenon-arc lamp (Lambda LS, Sutter Instruments) equipped with an excitation filter (470 nm CWL, 40 nm FWHM; Chroma ET470/40x) and directed toward the cranial window. Emitted light was collected through a tandem lens combination (Ratzlaff and Grinvald, 1991) consisting of a 55 mm lens and 85 mm lens affixed to the camera and passed through a longpass (495 nm cutoff, Chroma Q495lp) followed by a bandpass emission filter (525 nm CWL; 50 nm FWHM; Chroma HQ525/50). Images were acquired using StreamPix software (NorPix) controlling a CoolSNAP HQ2 CCD camera (Photometrics). After acquiring an image of the surface vasculature, the focal plane was advanced to a depth corresponding to ~300 – 400 μm below the brain surface. One trial of stimulation consisted of ~1 – 2 s of quiet, followed by sound onset (3 – 48 kHz frequency; 1 octave spacing; 1 s duration; 20 Hz modulation rate; 40, 60, 80 dB SPL) then 1 – 2 s of quiet. Each frequency-level combination was randomly repeated 5 times for a total of 75 iterations. Inter-trial interval was ~10 – 15 s. Acquisition of each frame was individually triggered and synchronized with the sound presentation using the Ephus software suite (http://www.ephus.org) (Suter et al., 2010).

### 2-photon imaging

To study cellular neuronal activity using 2-photon imaging, we used a scanning microscope (Bergamo II series, B248, Thorlabs) coupled to a pulsed femtosecond Ti:Sapphire 2-photon laser with dispersion compensation (Vision S, Coherent). The microscope was controlled by ThorImageLS software. The laser was tuned to a wavelength of λ = 940 nm in order to simultaneously excite GCaMP6s and mRuby2. Red and green signals were collected through a 16× 0.8 NA microscope objective (Nikon). Emitted photons were directed through 525/50-25 (green) and 607/70-25 (red) band pass filters onto GaAsP photomultiplier tubes. The field of view was 370×370 μm^2^. Imaging frames of 512×512 pixels (0.52 μm^2^ pixel size) were acquired at 30 Hz by bi-directional scanning of an 8 kHz resonant scanner. Beam turnarounds at the edges of the image were blanked with a Pockels cell. The average power for imaging in both L2/3 and L4 was <~70 mW, measured at the sample plane.

### Data Analysis

Wide-field image sequences were analyzed using custom routines written in Matlab (Mathworks). Images were parsed into trial-based epochs in which each frame sequence represented a single trial consisting of the presentation of a single sound frequency-intensity combination. For each trial, response amplitude (ΔF/F_0_) as a function of time was determined for each pixel using the formula ((F – F_0_)/ F_0_) where F corresponds to the time varying fluorescence signal at a given pixel and F_0_ was estimated by averaging the fluorescence values over 4 frames (~1 s) prior to sound onset for a given trial and pixel. For construction of sound-evoked response maps, the amplitude of the ΔF/F_0_ pixel response during 1 s after stimulus onset (~4 frames) was averaged across time and repetitions yielding an average response magnitude that was assigned to each pixel. Responsive areas in the average response maps were defined on a pixel-by-pixel basis as pixels in which the average brightness of the pixel during the 1 s after stimulus onset exceeded 2 standard deviations of the pixel brightness during the 1 s before the stimulus across stimulus repetitions.

### 2-photon image analysis

Image sequences were corrected for x-y drifts and movement artifacts using either the TurboReg in ImageJ (Thevenaz et al., 1998; Schindelin et al., 2012) or discrete Fourier transform registration (Guizar-Sicairos et al., 2008) implemented in Matlab (Mathworks) taking advantage of mRuby2 labeled neurons. Neurons were identified manually from the average image of the motion corrected sequence. Ring-like regions of interest (ROI) boundaries were drawn based on the method described in Chen et al. (Chen et al., 2013). Overlapping ROI pixels (due to closely juxtaposed neurons) were excluded from analysis. For each labeled neuron, a raw fluorescence signal over time (F) was extracted from the ROI overlying the soma. The mean fluorescence for each neuron was calculated across frames and converted to a relative fluorescence measure (ΔF/F_0_), where ΔF = (F – F_0_). F_0_ was estimated by using a sliding window that calculated the average fluorescence of points less than the 50^th^ percentile during the previous 10-second window (300 frames). Neuropil (NP) subtraction was performed on all soma ROIs (Peron et al., 2015). In short, the neuropil ROI was drawn based on the outer boundary of the soma ROI and extended from 1 pixel beyond the soma ROI outer boundary to 15 μm excluding any pixels assigned to neighboring somata. Thus, the final ΔF/F_0_ used for analysis was calculated as ΔF/F_0_ = (ΔF/F_0_)_soma_ – (α x (ΔF/F_0_)_NP_), where we used α = 0.9 (Peron et al., 2015) to reduce fluorescence contamination from the neuropil. Neurons in which the ΔF/F_0_ signal was significantly modulated by sound presentation were defined by ANOVA (p < 0.01) across baseline (pre-stimulus) and all sound presentation periods. Extraction of spikes from neuropil-corrected ΔF/F_0_ traces was performed using the Suite2P implementation of OASIS without an L1 penalty (Pachitariu, 2017). Suite2P generates a spike probability for each neuron in each imaging frame (λ). This continuous variable was then thresholded (λ_thr_) to yield a spike raster. Single neuron receptive fields (RF) were determined as the average ΔF/F_0_ response to each frequency-intensity combination across 5 stimulus repetitions during the stimulus window (1 s). Best frequency (BF) for each neuron was determined as the center of mass of the RF.

### Neuronal ensemble and avalanche analysis

Neuronal ensembles were defined based on contiguous frames in which at least 1 neuron was active (Figure 3b) following the approach by Bellay et al. (Bellay et al., 2015). Thus, ensembles were bounded by periods of silence in the population activity. Ensembles typically contained a varying number of active pyramidal neurons which could be active for only one or numerous frames within an ensemble. Ensemble size was defined as the number of neurons active summed over the number of frames each neuron was active. Ensemble duration, also called ensemble lifetime was defined as the number of frames constituting an ensemble. We note that similar to thresholding in electrophysiological recordings the specific value of λ_thr_ determines the number of spatiotemporal activity ensembles. If λ_thr_ is very low, most neurons are deemed active all the time, resulting in a small number of very large activity ensembles. Conversely, if λ_thr_ is very high, most neurons are considered inactive, again reducing the number of ensembles. Following the approach by Bellay (Bellay et al., 2015), we set λ_thr_ for each experiment to maximize the number of ensembles. Our results were not affected by small variations of λ_thr_ as demonstrated previously (Bellay et al., 2015). For analysis of evoked responses, ensembles were assigned to the temporal window when they initiated. Thus, an ensemble that initiated during the 1s stimulus window, even when it continued after stimulus offset, was categorized as an evoked ensemble.

To test if ensembles exhibit statistical properties of neuronal avalanches, we analyzed the probability distributions of both ensemble size and ensemble duration. First, the distributions for both size and duration were binned logarithmically, and probability density functions were obtained. For both the size and duration, we fit a power law or exponential distribution. We compared these fits and estimated parameters by calculating the log-likelihood ratio (LLR) by likelihood maximization as described previously (Clauset et al., 2009; Klaus et al., 2011). Slopes of the power law distributions (alpha values) were obtained using Kolmogorov-Smirnov goodness-of-fit to minimize the KS distance between the empirical distribution and a theoretical distribution for a range of alpha values (Clauset et al., 2009; Klaus et al., 2011). For controls, shuffled ensemble distributions were calculated from thresholded and shuffled spike density rasters using the same analysis outlined above. For shuffled rasters, each neuron separately had its spike probability estimates randomly permuted in time. This means the per-frame spike probabilities were maintained for each neuron, therefore maintaining the total number of active frames for each neuron. However, the individual temporal shuffling of neurons abolishes any temporal correlations between neurons. The shuffling process was repeated 10 times and the shuffled raster that produced the maximum number of ensembles was used.

Pairwise cross-correlation analysis was computed using zero-lag correlation of the λ estimates for all active neurons across the entire imaging session.

## Results

To compare responses of single neurons and neuronal ensembles, i.e. groups of groups, to auditory stimuli, we used *in vivo* 2-photon intracellular calcium imaging to sequentially measure activity of primarily pyramidal neurons expressing GCaMP6s in layer 2/3 (L2/3) and layer 4 (L4) in awake mice (n = 12 mice; Fig. 1a). Throughout the experiment the awake animal rested under the microscope while listening passively to semi-random presentation of 1 s short tonal sounds at 9 frequencies (3 – 48 kHz, two tones/octave) and 3 different sound pressure levels (40, 60, 80 dB) separated by 4 – 6 s. Each stimulus was repeated 5 times over the course of the experiment. We identified primary auditory cortex (A1) using widefield imaging (Fig. 1b). We then imaged the activity of neurons in L4 and L2/3 by imaging in different focal planes (Fig. 1c, d). For each neuron we extracted spike trains using standard methods (Pachitariu, 2017).

**Figure 1.**
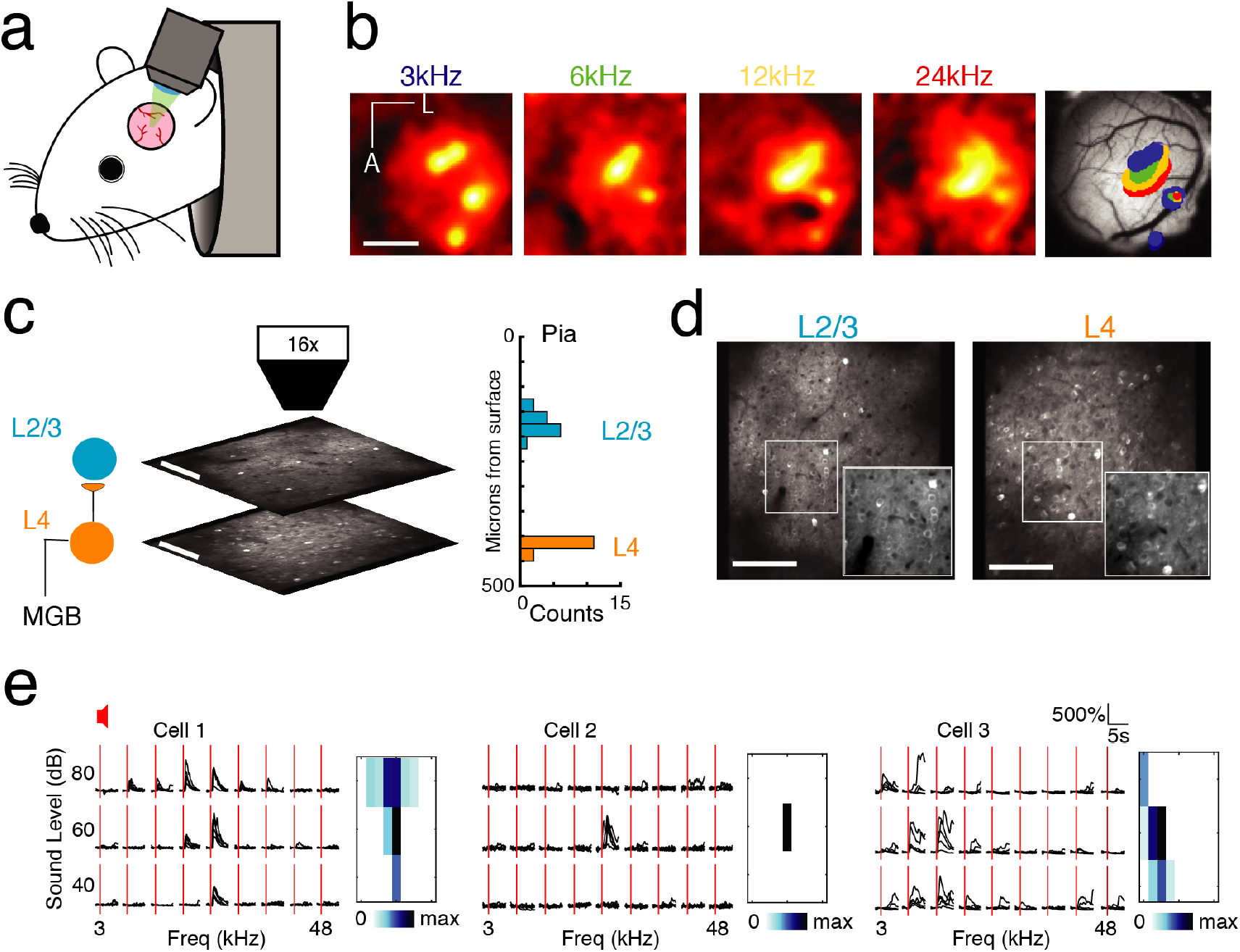
Imaging A1 population activity in the awake mouse. **a,** Cartoon of awake mouse under the microscope. **b,** Wide-field functional imaging identifies A1. Average activation maps from 4 different sound frequencies and thresholded activity overlaid on image of brain surface. Tonotopic gradient indicates location of A1. Scale bar = 1 mm. **c,** Imaging L4 and L2/3. Histogram shows distribution of imaging plane depths across all experiments. **d,** Representative fields of view from L2/3 (left) and L4 (right) showing GCaMP6s expression present in both layers. Scale bar: 100μm. **e,** Sound evoked intracellular calcium responses for 3 pyramidal neurons in A1 to 9 different frequencies (column) and 3 sound levels (row). Each black line indicates a single trial; 5 repeats per condition, red line indicates sound stimulus onset. *Right:* Frequency response areas.

First, we characterized single neuron receptive fields (RF) from the responses to single tones. Many neurons recorded in each layer were responsive to sound, i.e. showed an increase in firing during the stimulus period. Figure 1e shows three examples of RFs from A1 quantifying the change in stimulus specificity as a function of sound level. Stimulus specificity was found to broaden with an increase in sound volume (Fig. 1e, left), to exist only at a particular sound level (Fig. 1e, middle), or to broaden with a decrease in sound volume (Fig. 1e, right). Simultaneously imaged neurons in both layers showed heterogeneity of tuning in their responses (Fig. 1e) as reported previously in anesthetized and awake mice (Bandyopadhyay et al., 2010; Rothschild et al., 2010; Winkowski and Kanold, 2013; Maor et al., 2016; Liu et al., 2019).

### Population activity in A1 shows high variability

Given that observed local tuning heterogeneity of the imaged population is based on time-averaged measures, we sought to gain insight into statistical characteristics of the active neurons at a finer temporal scale. We first analyzed the properties of neuronal populations during ongoing activity in the absence of sound presentation. Ongoing activity could show periods of relative quiescence and periods of higher activity (Fig. 2a). On average ongoing activity was largely similar in both layers. Average firing rate was ~ 1.5 spikes/s (L4: 1.8 ± 1.11; L2/3: 1.83 ± 1.08; p>0.84). Firing in both layers was highly irregular as indicated by CVs between 1 – 2, with most irregular firing found in L4 neurons (Fig. 2a). Despite this irregularity, the distributions of pairwise cross-correlation were heavy-tailed with higher correlations found in L2/3 (Fig. 2c). These positive correlations indicate that neurons share common input and/or interact over distances during ongoing activity. These results are consistent with *in vitro* studies showing common inputs and connections in L2/3 of A1 (Oswald et al., 2009; Levy and Reyes, 2012; Watkins et al., 2014; Meng et al., 2017).

**Figure 2.**
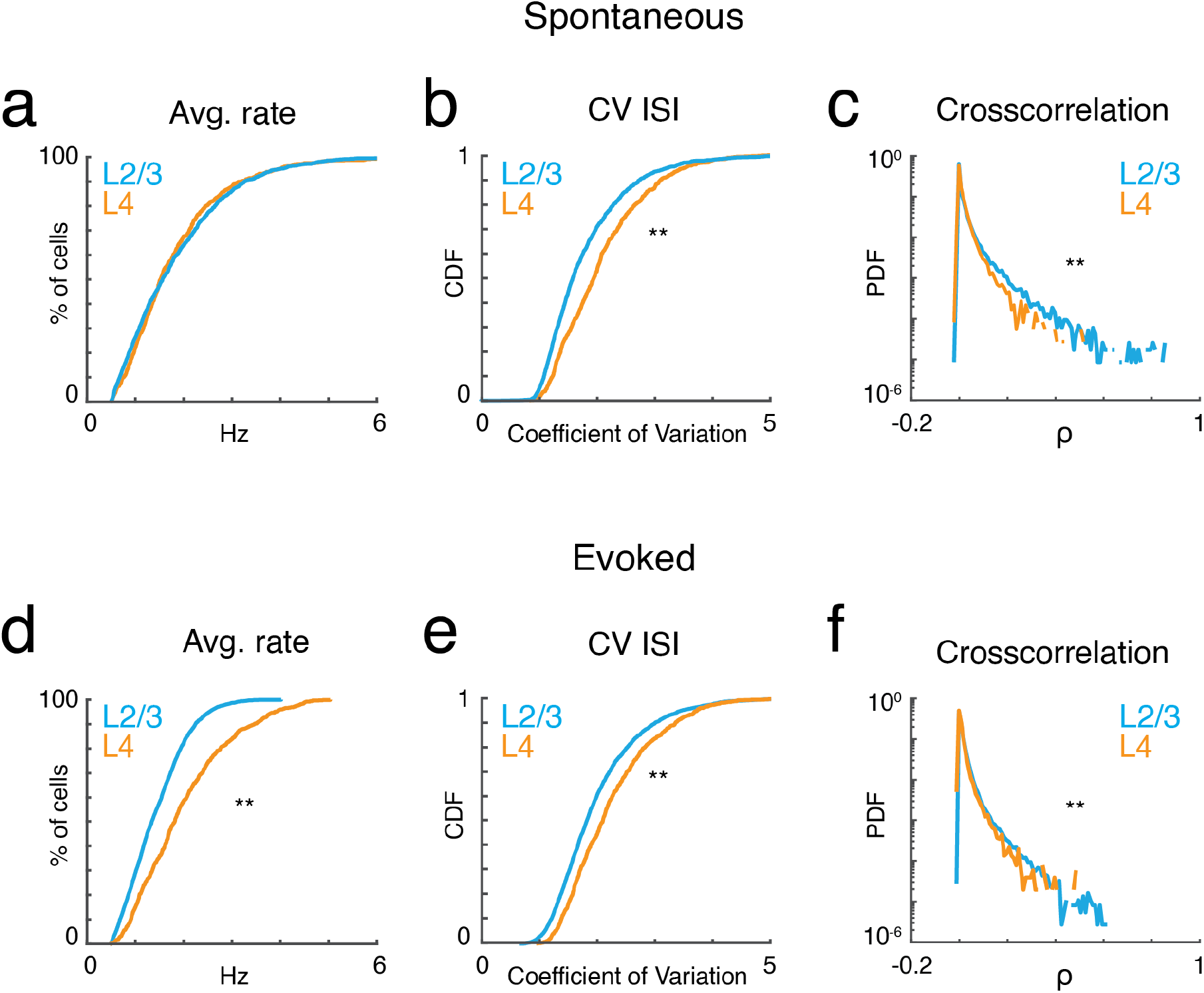
A1 activity is variable and correlated. **a,** Cumulative probability distributions of average spike rate for all recorded neurons in L2/3 (blue) and L4 (orange) during ongoing activity (p_rate_>0.84, ranksum). **b,** Corresponding cumulative probability distribution of inferred CV inter-spike interval (CV_ISI_) for all neurons in L4 (orange) and L2/3 (blue) (p_CV_<10^−20^, ranksum). **c,** Probability distribution of pairwise cross-correlation values for all neurons and experiments for L4 (orange) and L2/3 (blue) (p_CC_<10^−65^, ranksum). **d-f,** Average spike rate, CV_ISI_, and pairwise cross-correlation for evoked neuronal activity (p_rate_<10^−55^, p_CV_<10^−20^, p_CC_<10^−301^, ranksum).

Presence of a tonal stimulus increased firing rates and evoked rates were higher in L4 than L2/3 (Fig. 2d) consistent with electrophysiological recordings (Atencio and Schreiner, 2010). Evoked activity in both layers also showed high CV and heavy-tailed cross-correlations (Fig. 2e, f). The heavy-tailed nature of the correlation distribution suggests that despite the overall high variability observed during both ongoing and evoked activities, at least some neurons show relatively strong coordinated firing.

### Ongoing ensemble activity in A1 organizes as neuronal avalanches

To better understand the highly variable, yet seemingly coordinated organization of neuronal population activity, we grouped neurons based on the temporal association of their firing with other neuronal firings irrespective of spatial location.

We identified neuronal ensembles as groups of neurons active over successive imaging frames. Specifically, at a frame rate of 30 Hz, each frame defined a time bin of ~33 ms. Ensembles were bound by time bins that did not contain any active neuron, i.e. they exhibited successive time intervals in which at least 1 neuron was active (Fig. 3b). The total number of active neurons could vary between time bins with individual neurons becoming active and inactive, as long as contiguous population activity was maintained. Thus, ensembles could range from very small (a single neuron firing once within just one time bin) to large (many neurons firing over many successive time bins). To gain insight into the variability of ensembles, we quantified their sizes by summing the number of spikes (binarized) for each active neuron in each time bin. We then calculated the probability density function of ensemble sizes (Fig. 3c), for which qualitative inspection revealed a high probability of occurrence of small ensembles and a smaller probability of occurrence of very large ensembles. These probability distributions appeared linear in a logarithmic plot suggestive of a power law distribution. Indeed, the probability density distributions of ensemble size were significantly better fit by a power law compared to an exponential function (Fig. 3c,e; LLR_L4_ = 480.0 to 1251.9; LLR_L2/3_ = 551.8 to 1243.8; p<10^−13^; FOV_L4_ = 9; FOV_L2/3_ = 10). These power laws were abolished by temporal shuffling (Fig. 3c; LLR_L4-shuff_ = −690.9 to −175.1; LLR_L2/3-shuff_ = −424.9 to −108.8; p<10^−8^, favors exponential over power law) indicating that the temporal structure of ensemble activity is important in imparting the power law distribution of ensemble size.

**Figure 3.**
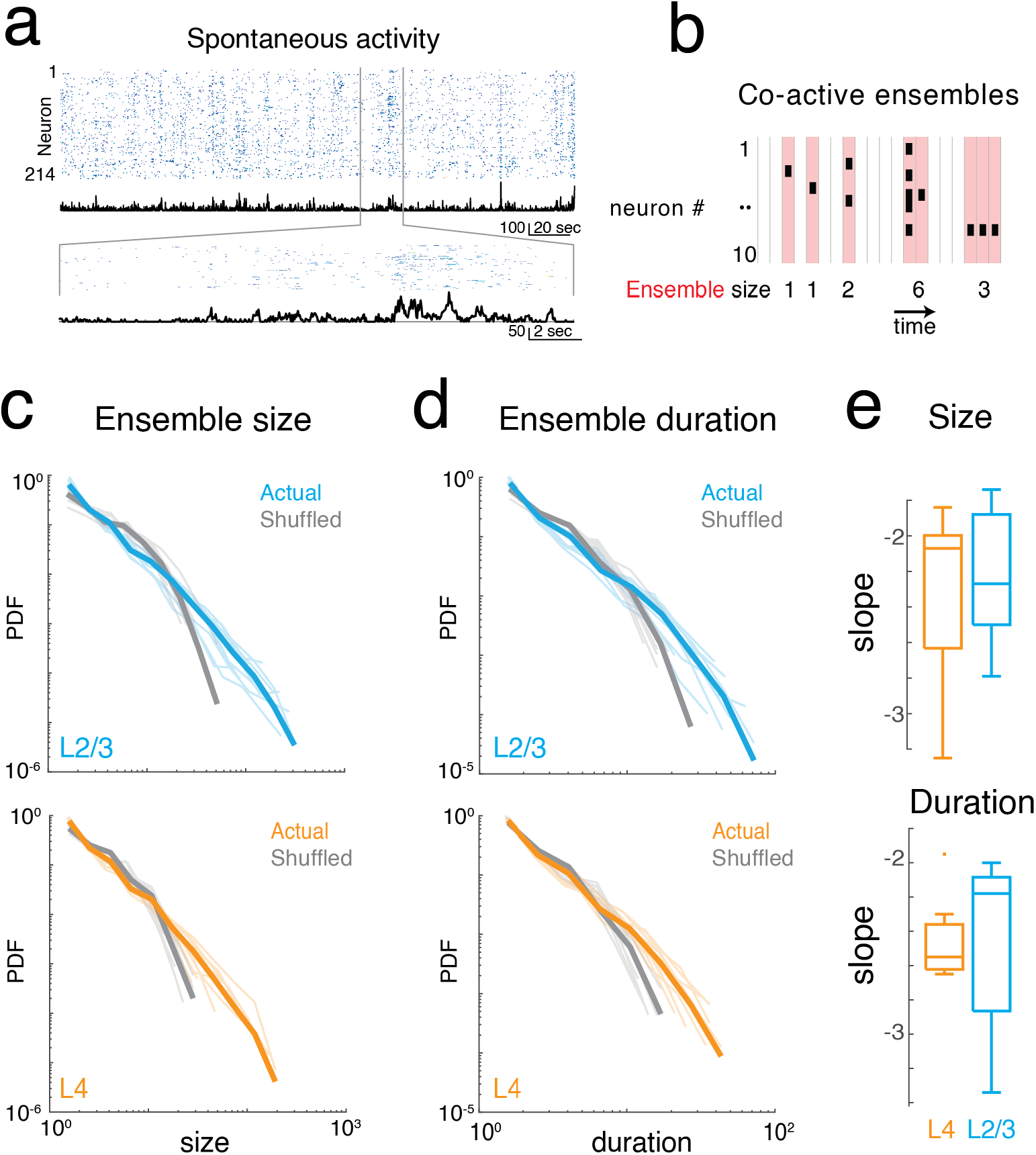
Imaging neuronal ensemble activity in awake mouse A1. **a,** *Top:* Binarized raster plot of ongoing activity recorded from L2/3 pyramidal neurons with corresponding population activities (black). *Middle:* Expanded view of raster plot revealing variable ongoing activity with population activities. **b,** Schematic depicting definition of activity ensembles from a neuronal activity raster. Activity from successive time bins/frames in which at least one neuron is firing (black rectangles) is concatenated into an activity ensemble (shading; 5 ensembles shown). Ensembles are separated by at least 1 frame with no activity. Vertical gray lines indicate boundaries of frame acquisition (Δt, 33 ms). Ensemble sizes defined as the number of spikes in each ensemble are indicated below. **c,** Probability density distributions of ensemble sizes from individual experiments in L4 and in L2/3. Gray lines indicate distributions from shuffled data. **d,** Probability distributions of activity ensemble duration of ongoing activity from all experiments in L2/3 and in L4. Gray lines indicate distributions from shuffled data. Faded lines indicate individual experiments. **e,** Boxplots showing slopes of probability distributions in ensemble size (*top*) and ensemble duration (*bottom*) for each individual experiment in L4 (orange) and L2/3 (blue), Size: α_L2/3_: −2.23 ± 0.39, α_L4_: −2.33 ± 0.52, Duration: α_L2/3_: −2.47 ± 0.49, α_L4_: −2.46 ± 0.26, mean±std p>0.65, unpaired two-sample t-test.

We next quantified the temporal duration of each ensemble by summing the number of consecutive time bins the ensemble was active. The distribution of ensemble duration also obeyed a power law distribution over an exponential (Fig. 3d,e; LLR_L4_ = 353.9 to 796.7; LLR_L2/3_ = 381.8 to 849.9; p<10^−34^), which was once again abolished by temporal shuffling (Fig. 3d; LLR_L4-shuff_ = 232.3 to 586.4; LLR_L2/3-shuff_ = 57.2 to 459.4; p<0.01, favors exponential over power law).

The presence of power-law distributions for ensemble size and duration suggests that ongoing ensembles in L4 and L2/3 organize as scale-invariant neuronal avalanches similar to ongoing activity in other cortical regions (Bellay et al., 2015).

### Evoked ensemble activity in A1 organizes as neuronal avalanches

We next investigated how the presence of a sound stimulus alters ensemble activity in A1. During an auditory stimulus, more A1 neurons fire and are thus recruited into ongoing active ensembles. Accordingly, sound evoked ensembles could deviate from a power law organization of avalanches by selectively increasing the incidence of large ensembles. Alternatively, sound evoked A1 ensemble activity could maintain scale-invariant organization by increasing the incidence of ensembles of all sizes.

From the event structure, we defined evoked ensembles as initiated during the 1s of sound presentation (Fig. 4). We then characterized the size and duration (lifetime) of the evoked ensembles by plotting the probability distributions of ensemble size and duration. Since varying sound levels potentially recruit different populations of neurons we separately analyzed ensemble activity at the three sound levels. All probability distributions appeared linear in a logarithmic plot suggestive of a power law distribution. Indeed, the probability density distributions of ensemble size for all sound levels were significantly better fit by a power law compared to an exponential function (Fig. 4b,d; Supp. Table 2). These power laws were abolished by temporal shuffling (Fig. 4b; Supp. Table 2; favors exponential over power law). We next quantified the temporal duration of each activity ensemble, which is the number of consecutive time bins for each ensemble. The distribution of ensemble durations also obeyed a power-law distribution (Fig. 4c,d; Supp. Table 2), which was once again abolished by temporal shuffling (Fig. 4c; Supp. Table 2, favors exponential over power law). The presence of power law distributions for ensemble size and duration suggests that evoked activity in L4 and L2/3 organizes as scale-invariant neuronal avalanches across different sound amplitudes. These results indicate that activity patterns are obeying the same statistical rules, however given the increase in activity of single neurons by sound stimulation it is expected that there are specific changes in the activity patterns due to the sensory stimulus. We thus calculated the average size and duration of evoked ensembles and found that both increased systematically with sound level (i.e., input strength) (Fig. 5a,b). Thus, while evoked ensembles in L4 and L2/3 remained power-law distributed in both size and duration (Fig. 4b,c, Supplemental Table 1, 2) the specific range from this distribution from which ensembles are chosen varies with sound level. Thus, scale-invariant avalanche dynamics in both layers L4 and L2/3 are preserved even in the presence of distinct sound input to A1 and across different sound amplitudes.

**Figure 4.**
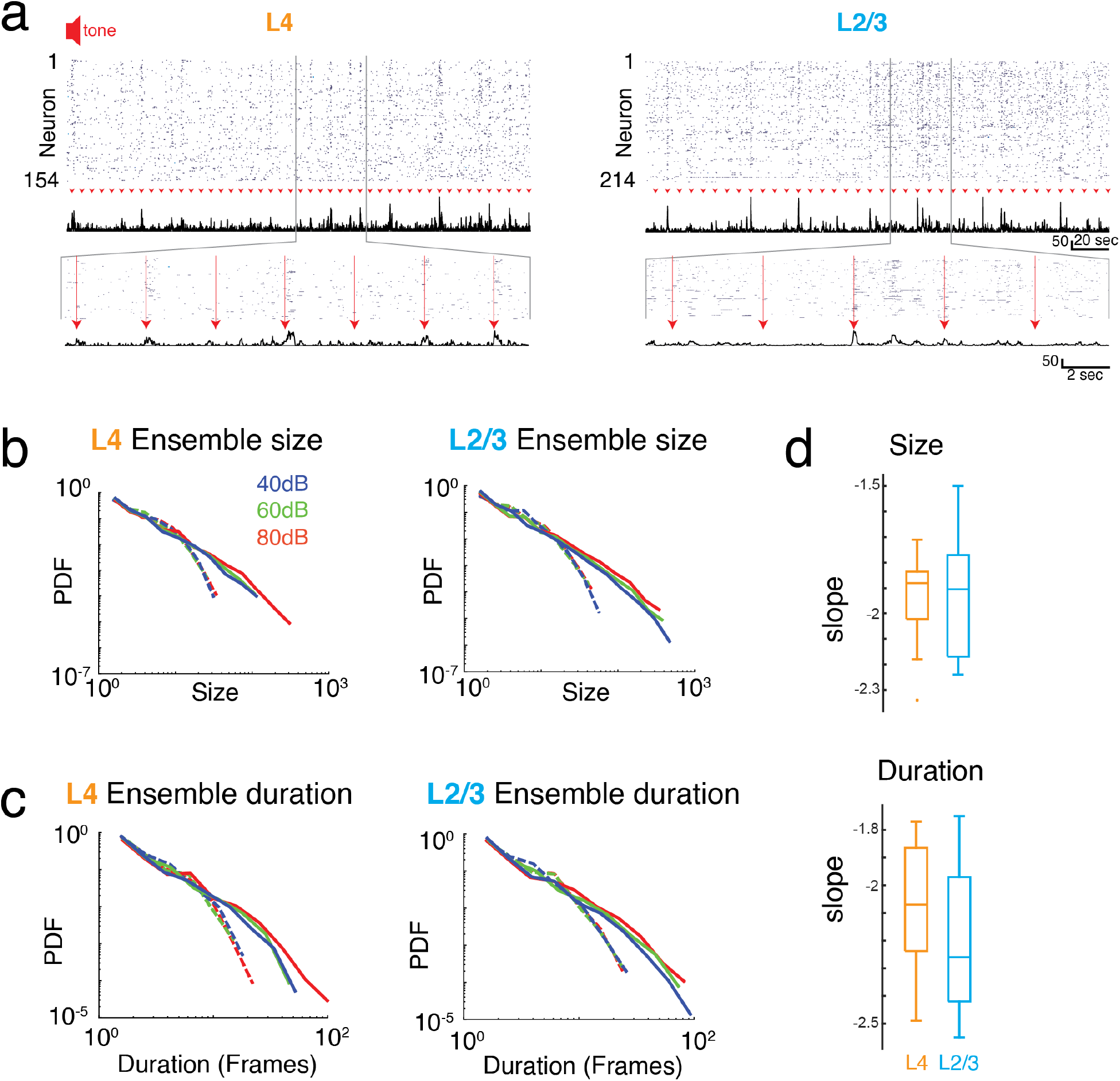
Neuronal ensembles in A1 during ongoing activity organize as neuronal avalanches. **a,** Activity raster of simultaneously recorded L4 (*left*) and L2/3 (*right*) neurons to brief auditory inputs (*arrow heads*). *Middle*: Corresponding population activities summed over all neurons. *Bottom*: Enlarged views revealing variable, transient evoked responses. **b, c,** Probability density distributions of ensemble sizes and ensemble duration reveal power laws for the evoked responses and different sound intensity in L4 (*left*) and L2/3 (*right*). Dashed lines indicate probability density distributions from shuffled data sets. **d,** Boxplots showing slopes of probability distributions in ensemble size (*top*) and ensemble duration (*bottom*) for each individual experiment in L4 (orange) and L2/3 (blue), Size: α_L2/3_:-1.93±0.25, α_L4_:-1.95±0.20, Duration: α_L2/3_:-2.20±0.26, α_L4_:-2.08±0.25, mean±std; p_size_>0.84, p_dur_>0.29, unpaired two-sample t-test.

**Figure 5.**
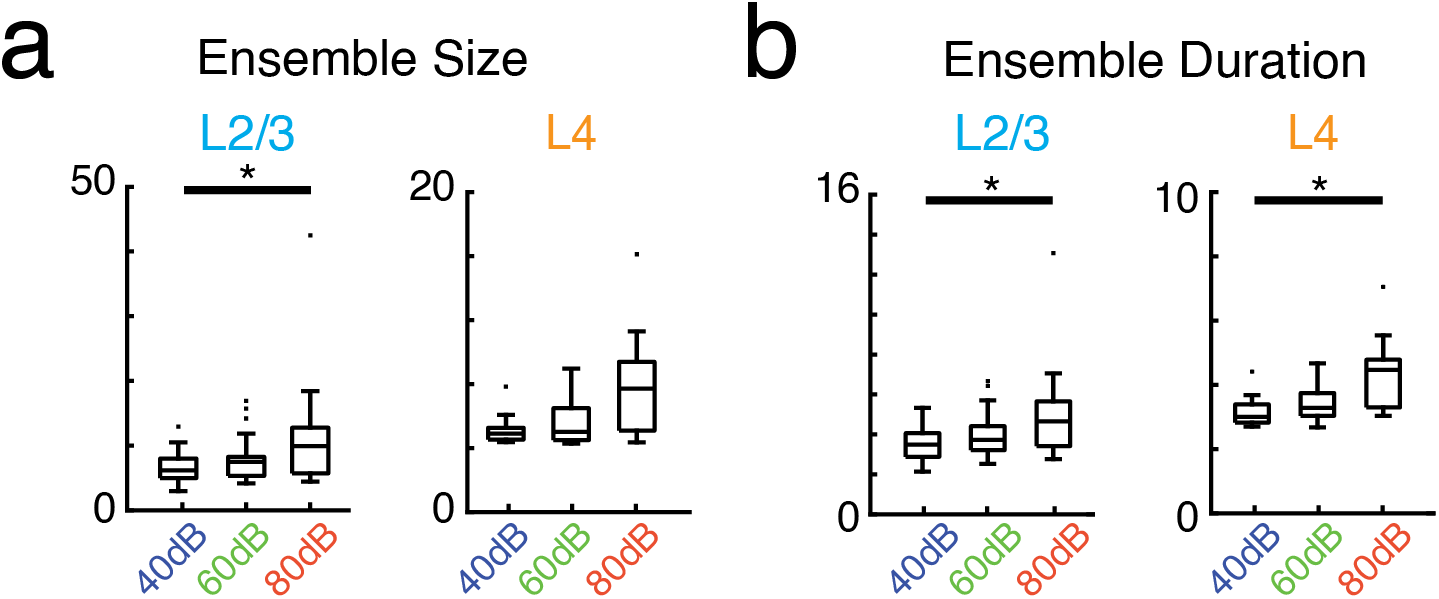
Average activity ensemble size and duration depend on sound level. **a,** Boxplots showing average ensemble size in L2/3 (left; p<0.04) and L4 (right) as a function of sound level. **b,** Conventions as in **a,** but for average ensemble duration (L2/3 p<0.04, L4 p<0.03).

### Evoked avalanches in A1 recruit neurons with widely varying tuning preference

Most behaviorally relevant sound stimuli are suprathreshold and thus recruit neurons with varying tuning preference (e.g. see Fig. 1e). Since we observe a wide range of ensemble sizes and duration we sought to determine if the range of frequency preference varied with ensemble size and duration. We thus quantified the tuning diversity in each evoked ensemble by measuring the interquartile range of BFs (IQR_BF_) from each active cell in the ensemble. We find that IQR_BF_ systematically increases with both ensemble size and duration (Fig. 6b, c) indicating that neurons of widely varying tuning preference are recruited into neuronal avalanches.

**Figure 6.**
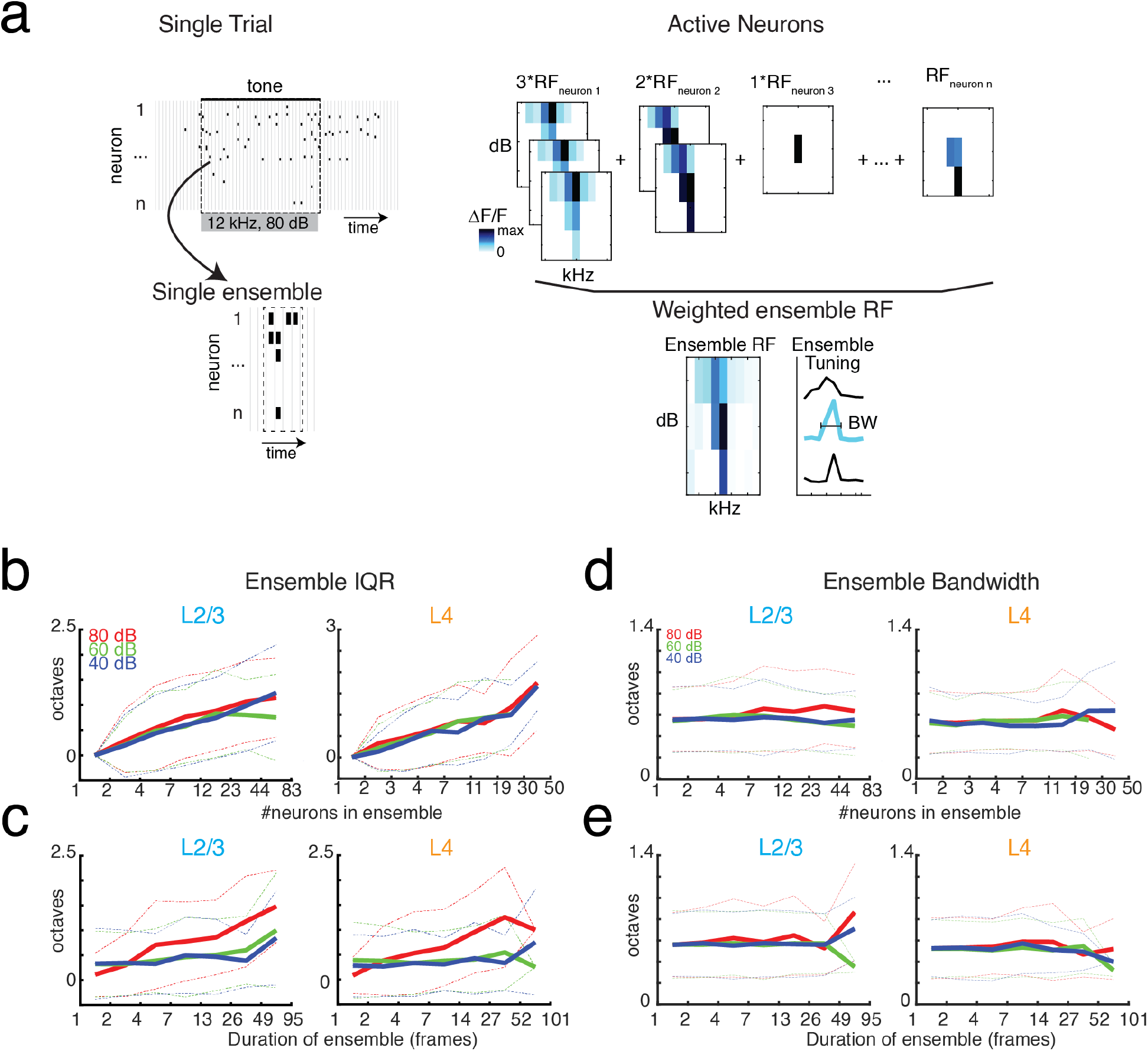
BF variability increases with ensemble size and duration, but bandwidth stays constant. **a,** Illustration showing the assembly of the ensemble RF from the weighted average of the RFs of individual neurons active in the ensemble. **b-c,** Ensemble IQR_BF_ as function of ensemble size (**b**) or duration (**c**). IQR_BF_ are binned according to logarithmically spaced bins of size and duration. Solid lines indicate mean. Dashed lines indicate one standard deviation. **d-e,** Conventions as in (**b-c**), but for ensemble bandwidth.

A neuron’s BF gives a limited description of a neuron’s overall receptive field. We thus also determined the compound receptive field of the evoked ensemble by averaging the receptive fields of the neurons active in the ensemble, weighted according to the amount of time each cell was active (Fig. 6a). The use of a weighted average is on the basis that the activity level of a neuron corresponds to its importance for encoding. We then determined the bandwidth of the ensemble receptive field. We find that ensemble bandwidth is not altered with respect to the size or duration of ensembles (Fig. 6d, e), indicating that stimulus selectivity is scale-invariant in the population activity. Furthermore, the bandwidth of the recruited ensemble did not vary with sound level indicating that ensemble receptive fields are sound level invariant, unlike many single-cell receptive fields.

## Discussion

We identified several fundamental properties of how A1 reliably represents stimuli in its spatiotemporal population response. First, we demonstrated that A1 represents even simple sound stimuli as temporally varying activity across distributed populations of diversely tuned pyramidal neurons. Second, pyramidal group activity in both layers obey similar statistical laws in that the size and duration of ongoing and sound evoked neuronal ensembles shows scale-invariance, the hallmark of neuronal avalanches. Finally, by introducing a novel approach of ensemble tuning, we found that neuronal ensembles recruit neurons of increasingly diverse tuning preference yet maintain constant selectivity with respect to ensemble size.

Our finding that both ongoing and evoked activities show signatures of neuronal avalanches is in line with experiment and theory associating avalanches with the optimization of numerous aspects of information processing such as dynamic range, information processing and transmission within layers (Shew et al., 2009; Shew et al., 2010; Shew et al., 2011; Shew and Plenz, 2013; Gautam et al., 2015) and between layers (West et al., 2008; Aquino et al., 2010; Aquino et al., 2011). Avalanche organization being maintained during stimulus presentation extends previous findings in *ex vivo* turtle visual cortex based on the local field potential demonstrating a return to avalanche organization after visual stimulation (Shew et al., 2015; Clawson et al., 2017). The existence of avalanche organization suggest that the network is poised to process incoming sensory stimuli with maximal dynamic range (Kinouchi and Copelli, 2006). The selective labeling of pyramidal groups in our experiments also specifically links avalanche dynamics to pyramidal groups, thereby extending previous reports on evoked avalanches in mice using a non-selective bulk labeling approach (Karimipanah et al., 2017). In summary, our work firmly suggests that A1 networks across layers and stimulus presentation optimize transmission of sound information by maintaining neuronal avalanche organization.

The reported statistics of neuronal ensembles are sensitive to sampling conditions. Spatial subsampling, in which only a small percentage of the population is measured, will change an expected power law distribution to an exponential form, an effect clearly visible at subsampling below 10% or more as shown in simulations (Priesemann et al., 2009; Ribeiro et al., 2010; Ribeiro et al., 2014; Levina and Priesemann, 2017). Subsampling might be the main reason why avalanches have not been observed in spike recordings using microelectrode arrays representing much lower subsampling conditions (Petermann et al., 2009; Ribeiro et al., 2010; Touboul and Destexhe, 2010). 2-photon imaging at cellular resolution allows for much higher neuronal sampling conditions and has been used previously to identify neuronal avalanches (Bellay et al., 2015; Seshadri et al., 2018) using non-negative deconvolution (Vogelstein et al., 2010). In the present study, we combined 2-photon imaging with an advanced, yet robust spike density estimation (Pachitariu et al., 2018) providing further support that spatiotemporal activity in spiking pyramidal groups organizes as neuronal avalanches. We also note that spiking between pyramidal neurons is significantly correlated, excluding models for the generation of power laws that feature no interaction between neurons (Martinello et al., 2017; Touboul and Destexhe, 2017). Our findings of robust power laws even under non-stationary conditions, i.e. evoked responses, are in line with previous reports in non-human primates based on the local field potential (Yu et al., 2017). We conclude that 2-photon imaging of cortical layers at cellular resolution in combination with robust spike density estimates allows for the clear identification of scale-invariant activity in ongoing and evoked population activity in primary auditory cortex.

We further demonstrate that avalanche organization and the population representation of stimulus information is largely independent of sound level. We introduce an ensemble RF measure that shows selectivity invariance with respect to stimulus sound level, in contrast to the widening of single-cell RFs across many auditory structures with increasing sound level (e.g. see Fig. 1e). This *in vivo* demonstration of a large dynamic range for A1 network activity matches reports on level-invariant perceptual discrimination performances (Jesteadt et al., 1977; Wier et al., 1977; Bernstein and Oxenham, 2006). It suggests that population coding of sound information might underlie level-invariant performances. Furthermore, we find that large activity ensembles recruit a diverse range of tuning preference, suggesting that BF does not fully represent a neuron’s role in population encoding.

In summary, our work reveals several key aspects of auditory cortical dynamics and in general cortical sensory coding. Even simple sensory stimuli are represented across distributed populations of diversely tuned neurons organized as scale-invariant spatiotemporal avalanches. We suggest that avalanche dynamics maximize information transmission between and within layers while providing an ordered framework for diversely tuned groups of neurons for which the relative timing of neuronal firing might be an important contributor to neuronal population encoding.

## Acknowledgements & Contributions

DEW, DP and POK designed research. DEW performed experiments. ZB, DEW, and SS analyzed the data. ZB, DEW, SS, DP, and POK discussed the results and wrote the manuscript. The authors thank Dr. Karel Svoboda for sharing designs enabling awake imaging and Dr. Mark Andermann for advice on awake imaging and Dr. Wolfgang Losert for helpful discussions. Supported by NINDS U01NS090569 (POK, DP), U19NS107464 (POK, DP), NIH RO1 DC009607 (POK), DOD W81XWH-16-1-0143 (POK, DEW), and the Division of Intramural Research of the NIMH ZIAMH002797 (DP). This study utilized the high-performance computational capabilities of the University of Maryland supercomputing resources (http://hpcc.umd.edu).

## Supplemental Tables

**Supplemental Table 1.** Statistical comparisons of the log-likelihood ratio values of avalanche size distributions and shuffled avalanche size distributions.

|  | L2/3<br>80 dB | L2/3<br>60 dB | L2/3<br>40 dB | L4<br>80 dB | L4<br>60 dB | L4<br>40 dB |
| --- | --- | --- | --- | --- | --- | --- |
| LLR Actual | 2573.9 | 2448.7 | 2094.4 | 1357.4 | 1480.3 | 1213.9 |
| Mean shuff | -0.3 | 14.2 | 3.6 | 104.3 | 99.8 | 115.5 |
| std shuff | 40.1 | 50.6 | 47.1 | 39.3 | 35.3 | 39.5 |
| Test | 't-test' | 't-test' | 't-test' | 't-test' | 't-test' | 't-test' |
| P-Value | $5.90 \times 10^{-181}$ | $1.40 \times 10^{-168}$ | $3.66 \times 10^{-165}$ | $7.05 \times 10^{-151}$ | $1.19 \times 10^{-159}$ | $4.52 \times 10^{-145}$ |

**Supplemental Table 2.** Statistical comparisons of the log-likelihood ratio values from avalanche duration distributions and shuffled avalanche duration distributions.

|  | L2/3<br>80 dB | L2/3<br>60 dB | L2/3<br>40 dB | L4<br>80 dB | L4<br>60 dB | L4<br>40 dB |
| --- | --- | --- | --- | --- | --- | --- |
| LLR Actual | 1361.4 | 1337.9 | 1246.8 | 824.0 | 946.8 | 817.5 |
| Mean shuff | 356.8 | 381.0 | 360.9 | 422.9 | 423.8 | 433.2 |
| std shuff | 36.6 | 41.5 | 42.0 | 35.3 | 36.6 | 35.9 |
| Test | 't-test' | 't-test' | 't-test' | 't-test' | 't-test' | 't-test' |
| P-Value | $1.56 \times 10^{-144}$ | $4.72 \times 10^{-137}$ | $3.25 \times 10^{-133}$ | $9.89 \times 10^{-107}$ | $1.75 \times 10^{-116}$ | $3.30 \times 10^{-104}$ |

## Notes

**Conflict of interest:** None

